# *Crossbrowse*: A versatile genome browser for visualizing comparative experimental data

**DOI:** 10.1101/272880

**Authors:** Sol Shenker, Eric Lai

## Abstract

The recent beyond-exponential growth in diverse collections of deep sequencing datasets creates enormous opportunities for discovery, concomitant with new challenges for displaying and interpreting these data. Notably, the availability of scores of whole genome sequences in multiple species clades enables comparative studies of functional elements. However, current genome browsers do not permit effective visualization of multigenome experimental data. Here, we present *CrossBrowse*, a standalone desktop application for displaying and browsing cross-species genomic datasets. We utilize data standards and graphic representation of popular browsers, and incorporate an intuitive graphical visualization of genome synteny that facilitates and drives human interrogation of comparative data. Our platform permits users with minimal informatics capacity to select arbitrary sets of genomes for display, upload and configure multiple datasets, and interact with vertebrate-sized genomic datasets in real-time. We illustrate the utility of *CrossBrowse* with interrogation of comparative invertebrate and mammalian datasets that provide insights into diverse aspects of transcriptional and post-transcriptional regulation. Of note, we show examplars of both preservation and divergence of functional elements that cannot be inferred from sequence alignments alone. Moreover, we demonstrate how inspection of primary data using *CrossBrowse* exposes an artifact in a typical strategy for assigning species-specific functional elements, and drives the implementation of an improved computational strategy. We anticipate that *CrossBrowse* will greatly foster user-based discovery within multispecies genomic datasets, and inform their bioinformatic interpretation.

## Introduction

Ongoing advances in next-generation sequencing (NGS) technologies, coupled with creative assays that interrogate diverse aspects of genomes and transcriptomes, are transforming the face of modern biology^1,2^. Indeed, many laboratories are now heavily reliant on techniques such as whole exome and whole genome sequencing, GWAS, ChIP-seq and genomic structure analyses, long and short RNA-seq, and so forth, all of which barely existed or were non-existent a decade ago. Many expedient software packages exist to process genomic and transcriptome data, and generate summary statistics of their properties. Nevertheless, human curation (i.e., “sequence gazing”) is still integral to effective and rigorous analysis. Not only is it prudent to provide reality checks regarding overall conclusions, the human eye is often instrumental in first recognition of novel genomic phenomena that are not appropriately handled by existing software packages. In turn, such insights can fuel subsequent computational work.

As an example from our experience, while transcripts were annotated through a standard pipeline in the modENCODE project^3^, our visual recognition of long, neural 3’ UTR extensions that were not assembled as continuous exons led us to perform the initial annotation of the transcriptome via end-to-end manual browsing of the *Drosophila* genome^4^. The broad foundation we gained by human curation was critical to develop our de novo transcriptome assembly and analysis method IsoSCM^5^, which enables effective analysis of alternative polyadenylation trends from RNA-seq data. Thus, effective data visualization can facilitate human-driven identification of novel genomic patterns that would be otherwise overlooked by existing analysis tools, and guide the development of new analytical strategies.

The burgeoning amount of comparative genomic datasets creates new challenges in genomics visualization. One of the most-widely used public genomic resources is the UCSC Genome Browser, a portal for real-time user interactions with multiple genome alignments. In addition, a multitude of genomewide datasets are publicly hosted by the UCSC Genome Browser, and one can also link to private datasets. However, a clear limitation is that the UCSC platform does not permit comparison of experimental data from multiple species, nor visualization of genome rearrangements beyond linear relationships. While the UCSC comparative assembly hub introduce snake tracks to represent structural genome changes^6^, substantial issues remain to navigate and interpret multi-genome datasets.

The democratization of next-generation sequencing means that it is now straightforward to generate genomewide datasets in historically non-model organisms, and public deposits of such cross-species data now present largely untapped opportunities for discovery. Moreover, the advent of CRISPR/Cas9 genome engineering now means that a much broader range of species are experimentally and genetically tractable. Thus, genomics data can not only help interpret the genomes of “classic” model organisms, but serve as the basis for experimental designs in new species. Nevertheless, while cross-species datasets provide rich resources for bioinformatics investigations, they are currently largely inaccessible to biologists and laboratories lacking significant computational expertise.

While some approaches have begun to address this problem, none of the available cross-species browsers satisfactorily deal with the challenges of current genomics data. Combo was one of the first comparative browsers^7^, but it does not handle NGS data or permit more than two genomes to be displayed. Sybil^8^ handles only bacterial-sized genomes, displays only annotated genes, and does not support NGS formats. mGSV^9^ is a web-based browser through which one must upload data prior to visualization, and is consequently slow and not secure. In addition, it limits the user to loading only one data track, and uses a custom upload format that can render data configuration tedious. Finally, GBrowse_syn^10^ requires a dedicated Linux server, and significant command line usage to modify session configuration, limiting breadth of its utilization. In addition, all of these packages share the significant limitation of showing only a fixed coarse-grain representation of synteny relationships, and thus lack dynamic visualization as the user zooms between multi-gene and the nucleotide level resolutions. This is a signature feature of the UCSC Genome Browser and the Integrated Genome Viewer that today’s biologists have come to expect. Reciprocally, the UCSC Genome Browser focuses on local, linear alignment relationships, at the expense of visualizing interrupted segments (indels) and larger genomic rearrangements. Finally, while web-based genome browsers have advantages with respect to aggregating datasets and a means to share a session among collaborators, this design can place inordinate amount of strain on server, degrading responsiveness and user experience.

We sought to address all of these limitations in *CrossBrowse*, our standalone desktop application for cross-species data visualization for vertebrate-sized genomes. To create a tool familiar to users of existing genome browsers, we took advantage of data standards and graphical representation established by first generation genome browsers, and combined it with an intuitive graphical representation of genome synteny relationships and multispecies experimental data. We demonstrate the utilities of *CrossBrowse* for analyzing transcriptional and post-transcriptional regulation, and highlight its attributes and capacities that permit bench scientists to interact with and make discoveries from cross-species genomics data.

## Results and Discussion

### A genome browser with adaptive granularity visualization across syntenic loci

We designed a browser to meet the aforementioned challenges for multi-species visualization of aligned genomes and experimental datasets. Here, we describe key features of *CrossBrowse*, especially with regard to issues not satisfactorily handled by existing browsers.

*CrossBrowse* generates the synteny visualization using whole genome sequence alignments. These allow *CrossBrowse* to translate coordinate positions between reference genomes, the same functionality provided by the *liftOver* tool from the UCSC genome browser. *CrossBrowse* reads whole genome alignments from CHAIN-format files that can be downloaded from the UCSC genome browser. The window is composed of multiple stacked “traditional” genome browsers, which are connected by graphics that illustrate syntenic relationships. These are represented as colored “blocks” that indicate boundaries of homologous regions. Unaligned sequences are left white, but the coloring scheme easily allows one to identify when such regions reside within a larger syntenic region. When sequences reside on opposite strands with respect to the reference genome, the block will reflect this inversion.

Since there can be extensive homology between related genomes, *CrossBrowse* divides each longer block of homology into a series of smaller blocks, so that the synteny visualization adapts to the viewed coordinates. The synteny visualization reacts to user interaction as the mouse moves over the view area, allowing for the rapid assessment of information at orthologous positions, even in the context of structural genome changes (**Supplementary Video 1**). Syntenic regions are shaded to facilitate the location of orthologous positions across genomes. The coloring scheme is stable with respect to the displayed coordinates, thereby maximizing visual continuity as the view is adjusted. We note that tools such as Sybil and mGSV and Gbrowse_syn generate conceptually similar graphical motifs to visualize synteny. However, only *CrossBrowse* can represent synteny with a granularity that adapts to the user’s view. Thus, *CrossBrowse* can be readily be applied to investigate features ranging from megabase windows all the way down to single nucleotide resolution. A video that illustrates the adaptive granularity of the synteny visualization is provided as **Supplementary Video 2**.

We illustrate the utility of synteny visualization using homeobox gene re-arrangements observed in the *Drosophila Antennapedia*-Complex. Hox genes are essential to establish segmental identities during development and exhibit striking correspondence between their relative expression domains along the anterior-posterior axis and their linear order along the genome. The linear order of Hox complexes is generally conserved, but at least seven structural re-arrangements are observed across the Drosophilid phylogeny^11^. Such re-arrangements are not evident when using popular interfaces such as the UCSC genome browser or IGV, and are often rendered manually (**Figure 1A**). However, using *CrossBrowse*, one can easily zoom in and out of this complex genomic region and observe the phylogenetic relationships of their changing gene orientations in a graphically intuitive manner (**Figure 1B**).

**Figure 1:**
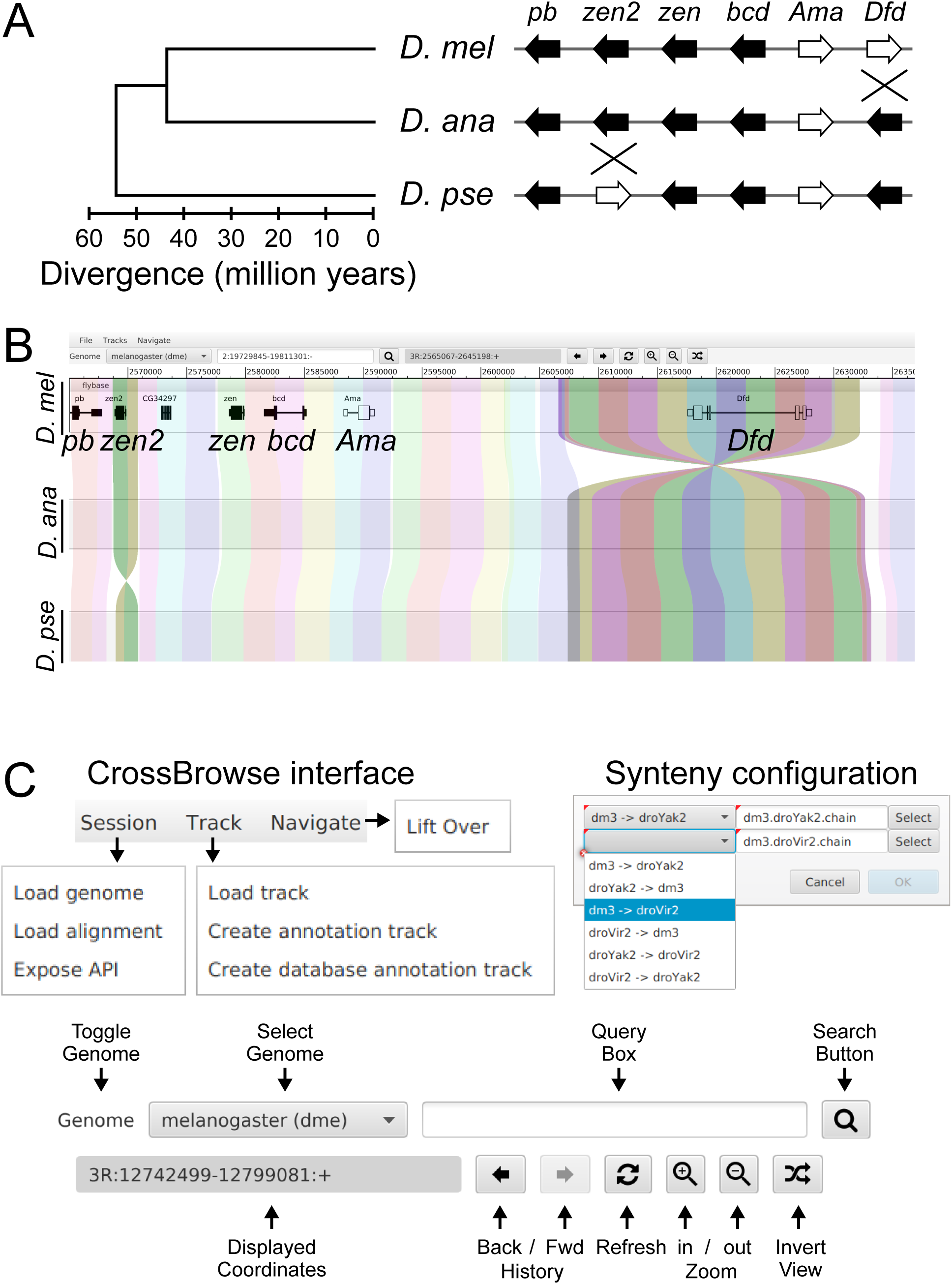
Structural changes across Hox complexes visualized using *CrossBrowse*. (A)Graphical representation of structural changes within the *Antennapedia*-Complex locus across the radiation of *D. melanogaster, D. ananassae*, and *D. pseudoobscura*. The separation of *D. pseudoobscura* from *D. melanogaster* and *D. ananassae* is accompanied by inversion of *zen2*. Likewise, the speciation of *D. melanogaster* and *D. pseudoobscura* is accompanied by an additional inversion of the *Dfd* locus. (B) A *CrossBrowse* screenshot illustrates how these structural re-arrangements occurring at the *Antennapedia*-Complex are readily visualized using *CrossBrowse*. A track displaying gene structure is displayed in the melanogaster genome. The location of syntenic genome sequences are indicated by ribbons that span the individual genome views. Structural inversions relative to neighboring genomes are represented as twists in the ribbon. Darker shading is used to highlight the location of the structural changes. (C) Configuration of the browser is easily achieved using a graphical interface. Below, the navigation bar is annotated with the function of each button. The popup at right demonstrates the procedure for loading whole genome alignment CHAIN files.

### An intuitive and facile interface for concurrent exploration of multiple genomes

*CrossBrowse* implements navigation features and graphical standards established by existing browsers, thereby providing a familiar user environment. As seen in the example *CrossBrowse* session (**Figure 1C**), the top of the interface displays *Session, Track*, and *Navigation* menus. The *Session* menu provides an interface for the user to configure which genomes are displayed in the browser, and pairwise alignments between them. The *Track* menu is used to load tracks in the browser, and adjust how they are displayed. The *Track View* is partitioned into separated stacks of *Track*s when the *CrossBrowse* session is configured to display multiple genomes. The *Navigation* menu provides access to functionality for selecting views of syntenic regions between reference sequences, access navigation history, refresh tracks, and adjust the zoom level.

We incorporated features that facilitate genome browsing in the context of multiple genomes. For example, coordinate conversion is tightly integrated with the browser, and is accessed through the toolbar by selecting *Navigate*→*liftOver*. The source genome for the *liftOver* can be selected from the drop down menu at the top left, and the list of target regions identified are displayed in the list below. Selecting one of the target regions and pressing “*liftOver*” will set the coordinates in the target genome based on the region of synteny with the window displayed in the source genome (**Figure 1C**).

One issue with CHAIN files is that the existing tools for coordinate conversion (which are necessary to build the overlay visualization) are designed for batch queries. Given a set of intervals in one genome they read the entire CHAIN file, from start to finish, and report the translated coordinates for all query segments that can be mapped to the target genomes. This process is too slow to support interactive cross-species data visualization, since the delays in refreshing each view of adjusted genome coordinates are unacceptable for browsing data. To enable low latency coordinate conversion, *CrossBrowse* uses TABIX indexing^12^ to accelerate access to CHAIN file genome alignments. The UCSC genome browser hosts CHAIN files generated with respect to a popular reference organisms (e.g., relative to human, mouse, fly). However, bidirectional coordinate conversion requires these alignments to be available in both directions, which would require two CHAIN files. To simplify configuration and allow users to easily use publicly available CHAIN files, *CrossBrowse* includes the functionality for deriving inverted CHAINs, obviating the need for the user to perform unnecessary pre-processing.

A further limitation of existing tools for multi-species visualization is the difficulty for configuration. While the session for many popular single genome browsers can be conveniently configured via an intuitive graphical user interface, existing multi-species browsers require the user to install the browser on a web server. These browsers require substantial command line usage to configure and install the browser (GBrowse_syn^10^) and/or convert data into browser-specific formats (mGSV^9^). In contrast, *CrossBrowse* provides a graphical interface, making the browser more readily accessible to users lacking the time or technical expertise to install and configure a session. The only requirement for running *CrossBrowse* is a Java virtual machine (JVM) which is readily available on all operating systems.

Upon launch, the user is presented with an empty browser session, within which a user can load genomes, CHAIN files, and data tracks from the menu bar. Once configured, the synteny display is synchronized as the user navigates directly to a specific genomic coordinate or using text search against loaded annotations. A detailed text description that illustrates the process of configuring and navigating a multi-species session is provided as **Supplementary File 1**.

### Practical applications of *CrossBrowse*

Coding sequences are often broadly conserved, but strong constraints on other genomic elements can be used to infer their underlying functionality and utility (e.g., TF binding sites, miRNA and RBP binding sites, etc.). While existing browsers (e.g. UCSC genome browser) permit facile visualization of conserved sequences alongside experimental data from one organism, they do not permit simultaneous visualization of multispecies data. This is needed to distinguish whether utilization of a conserved functional element is similar, or perhaps divergent between species. A greater challenge is that many genomic elements are under modest positional constraint and/or difficult to infer directly from primary genome sequence alone. Thus, it can be difficult to distinguish functional elements that are functionally conserved but are defined by different primary sequences, from those that are truly species-specific and evolutionarily divergent. We illustrate the general utility of *CrossBrowse* with analyses of invertebrate and mammalian genomic data, which demonstrate insights into diverse aspects of transcriptional and post-transcriptional regulation.

### Evolutionary dynamics of enhancers and insulators

Facile browsing provides any bench scientist the ability to evaluate general trends in genomewide comparative data, as well as to navigate specifically to genes of particular interest for functional study. To illustrate the capability of *CrossBrowse* to handle mammalian-scale datasets, we downloaded ChIP-seq data for several chromatin features (H3K27ac, P300, TFAP2A) used to identify divergent enhancers between human and chimp^13^. By navigating to the *COL13A1* locus, the resulting visualization readily illustrates the human-biased enhancer noted in the prior work (**Figure 2A**, red box). Moreover, one can identify a weakly human-biased enhancer (yellow box) as well as a strongly chimp-biased enhancer (blue box). This emphasizes the ease with which *CrossBrowse* can be configured for retrospective analysis of published datasets.

**Figure 2:**
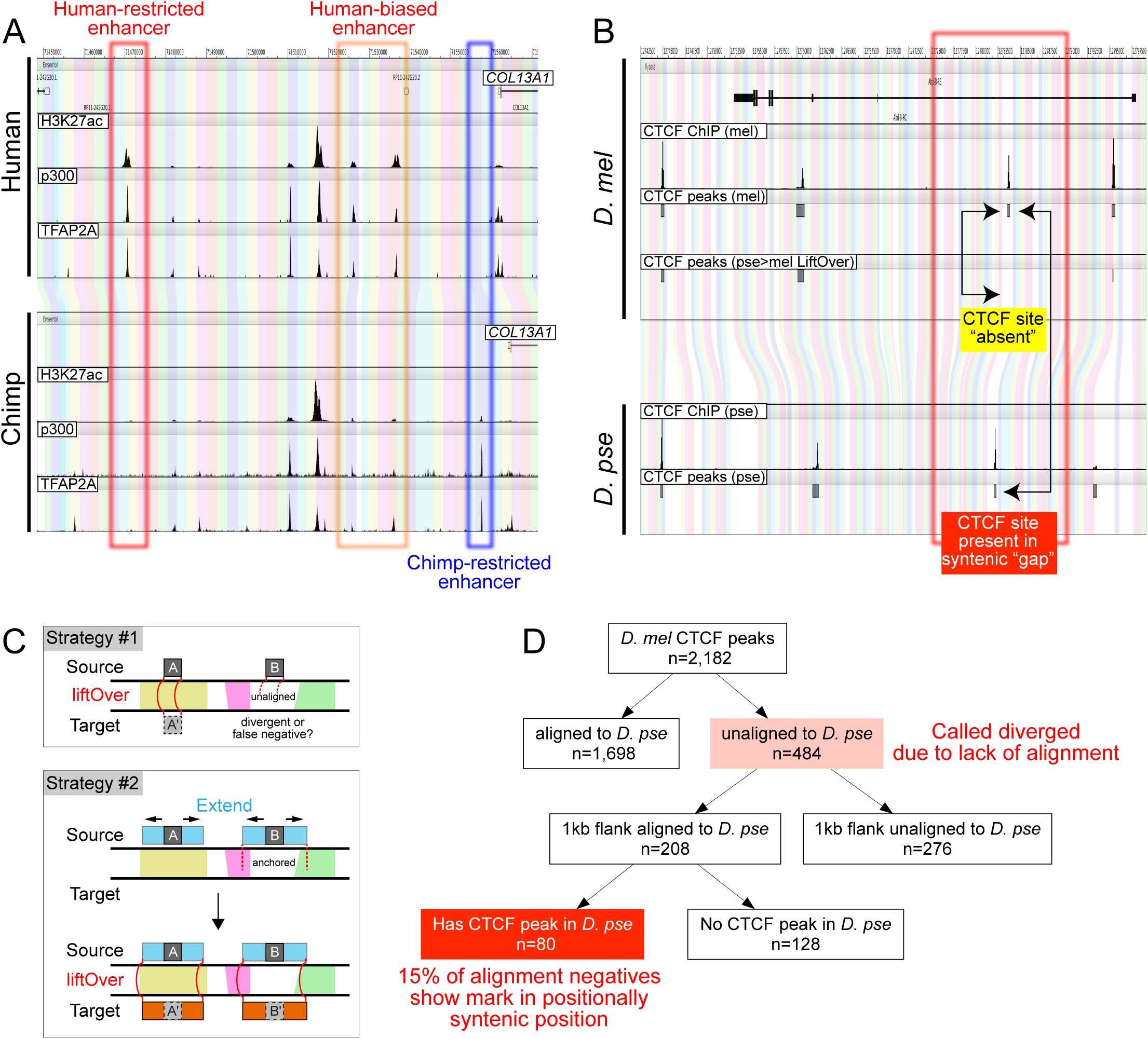
Visualization of cross-species chromatin-based data using *CrossBrowse*. (A) Screenshot of a *CrossBrowse* session that allows interrogation of species-specific enhancer evolution at mammalian *COL13A1* reported by Wysocka and colleagues^13^. Here, we plot WIG tracks of their H3K27ac, P300, and TFAP2A ChIP-seq data from human and chimp. The red box highlights the human specific enhancer earlier reported^13^, and the blue arrow indicates the presence of a chimp biased enhancer. (B) Screenshot of a *CrossBrowse* session comparing CTCF sites assayed across multiple Drosophilid species. The *Abd-B* locus was originally reported to harbor evolutionary divergent CTCF binding by White and colleagues^16^. The red box highlights a CTCF binding in *D. melanogaster* that is not directly aligned in *D. pseudoobscura*, and is thus lost in the standard *liftOver* translation. However, visual affirmation of their synteny is easily established via proximal flanking regions. (C) Comparison of strategies to ascertain conservation and divergence of functional elements. Strategy #1 was used in the original CTCF study^16^, and is typically employed in cross-species studies. ChIP-seq peak coordinates were directly translated using *liftOver* when a sequence alignment is available (element “A”, left), while peaks residing within segments that are locally unaligned are discarded (element “B”, right). As informed by *CrossBrowse* visualization, we implemented the modifications shown in Strategy #2. First, CTCF peaks were extended by a fixed distance (500 bp) on either side (blue rectangles). If extended peaks could be anchored in flanking aligned regions, coordinate translation was performed to identify homologous regions (red connectors). This strategy allows one to query both elements A and B for functional conservation in the experimental data. (D) Flowchart for re-analysis of comparative CTCF ChIP-seq data^16^. Of 484 events in *D. melanogaster* originally called as absent in *D. pseudoobscura*, due to lack of *liftOver* coordinates, we were able to rescue nearly half of them as residing in clearly orthologous regions based on their genomic flanks. About 40% of these regions (comprising 16.5% of total alignment negatives in the *D. melanogaster →D. pseudoobscura liftOver*), indeed harbored conserved CTCF binding in the latter species. We vetted all 80 locations by manual inspection in *CrossBrowse*.

In comparative genomic studies, it is typical to designate one genome as a reference, and translate the coordinates of all measurements from all experiments made in non-reference genomes into a common coordinate system of the *reference* genome. This “coordinate translation” between regions with sequence similarity is performed using tools such as *liftOver* and *CrossMap*^14,15^ using pre-calculated whole genome alignments. While this approach works well for comparing features that occur in well-aligned and clearly orthologous regions, care must be taken to design the analysis to properly account for events that occur in regions without an easily identified homologous sequence. Although such events are typically considered in bulk to be species-specific, the direct analysis of coordinate translated measurements can create artifacts of interpretation.

We illustrate this case using comparative data for the insulator DNA-binding protein CTCF, analyzed across 4 *Drosophila* species^16^. This work identified conserved-and species-specific CTCF binding events, and concluded that while CTCF DNA binding motif is conserved, the location of binding events between species evolves dynamically. In particular, *liftOver* coordinate-translation of CTCF ChIP-seq data at *Abd-B* was used to identify cases of CTCF binding in *D. melanogaster* that were lost in *D. pseudoobscura*^16^ (**Figure 2B**, yellow box). However, comparative visualization of the CTCF ChIP-seq data aligned to each cognate genome in *CrossBrowse* reveals that CTCF binds at a similar genomic position in both species (**Figure 2B**, red box). This region is anchored by flanking syntenic sequences, even though the CTCF binding itself occurs within a region that was not well-aligned. This situation can be classified as a “false-positive” from the *liftOver* pipeline, since CTCF binds in an analogous genomic position in these species relative to the genes its insulator activity should affect.

Given that coordinate translation of sequencing data can introduce such false-positive calls, we were interested to assess its impact genomewide, and to develop a refined analysis strategy. In particular, we sought to distinguish true species-specific events from ones that occur in regions of locally low sequence identity, whose overall genomic correspondence was anchored by nearby flanking orthologous regions (as observed in the *Abd-B* locus). To do so, we expanded the binding peak called from the ChIP-seq data by 500bp on each side before performing coordinate translation (**Figure 2C**). Using this strategy we were able to identify >200 CTCF binding events in *D. melanogaster* that failed standard *liftOver*, but where the flanking genomic DNA can be used to identify the syntenic position in the *D. pseudoobscura* genome (**Figure 2D**). Of these, browsing confirmed CTCF binding within the syntenic segment in ~40% of the “rescued” genomic regions (80 loci, comprising 15% of the total CTCF peaks).

We emphasize that manual validation is extremely cumbersome using existing browser solutions; i.e., necessitating the identification of orthologous regions and navigating to the respective data in separate browsers. These maneuvers are sufficiently onerous that they usually preclude systematic manual assessment of species-specific features. Our detection of a substantial number of falsely positive functional divergences highlights how visual browsing of multispecies genomic data in *CrossBrowse* can inform computational approaches for improved comparative analyses.

### Evolutionary dynamics of alternative polyadenylation

In addition to visualizing the evolution of DNA-based functional elements, *CrossBrowse* has diverse utilities for analysis of transcriptome elements. RNA-seq has been used to study the evolutionary conservation of alternative splice isoforms^17^-^19^ and alternative polyadenylation (APA) events^5,20^. We previously described tissue-specific utilization of 3’ UTR isoforms^4,21^, including broad trends for expression of shorter 3’ UTR isoforms in testis and longer 3’ UTR isoforms in neural tissues.

Despite the qualitative similarity of these tissue-specific patterns, proximal polyadenylation sites often lack clear signatures for primary sequence conservation, as we observed in *Drosophila*^4^. To better understand how the observed primary sequence contributes to alternative isoform expression between species, we analyzed RNA-seq and 3’-seq data from head and testis of *D. yakuba*, which is only ~6 million years diverged from *D. melanogaster* (Sanfilippo and Shenker et al, in preparation). Examination of differential isoform expression patterns using *CrossBrowse* reveals different modes of 3’ UTR expression pattern evolution between species.

In **Figure 3A**, the *CG2201* locus provides an example where loss of the testis-specific polyadenylation in *D. yakuba* results in loss of highly differential 3’ UTR expression between tissues observed in *D. melanogaster*. We find it remarkable to find such divergent mRNA processing patterns over recent evolutionary history. By contrast, *Pde1c* illustrates a case where despite the fact that both fly species exhibit 3’ UTR shortening in testis relative to head, the actual sites of proximal polyadenylation are not at orthologous positions (**Figure 3B**). This type of “divergent, yet functionally similar” gene regulation (akin to the CTCF example, **Figure 2B**) would have been difficult to appreciate without simultaneous browsing of data mapped to each cognate genome.

**Figure 3:**
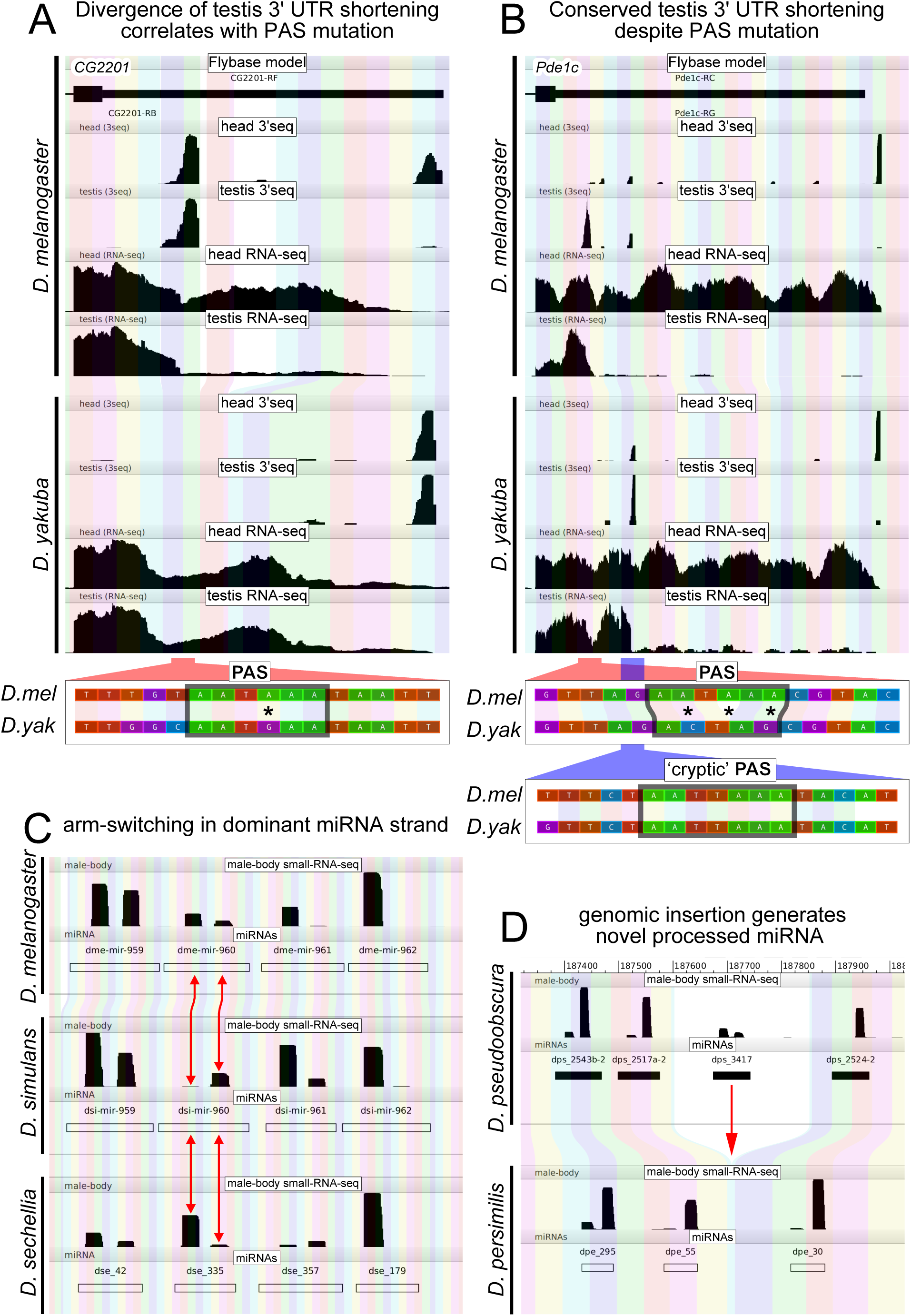
Visualization of cross-species transcriptome-based data using *CrossBrowse*. Shown are examples of how diverse types of transcriptome data can be interpreted using *CrossBrowse* to identify evolutionary divergence in functional elements. (A, B) Screenshots of a *CrossBrowse* session loaded with total RNA-seq and 3’-seq data from heads and testes of *D. melanogaster* and *D. yakuba*; note that 3’ UTRs are not annotated in existing *D. yakuba* gene models. These panels illustrate loci utilizing tissue-specific alternative polyadenylation that are evolving in distinct manners between these species. (A) Top, in the case of *CG2201*, the proximal testis PAS is lost in *D. yakuba*. Below, zooming to the nucleotide level shows that the canonical testis PAS used in *D. melanogaster* is diverged in *D. yakuba* (red box, asterisk). (B) The higher level view at *Pde1c* shows that both species incur testis-specific 3’ UTR shortening, but the color-coded alignments indicate that non-orthologous termini are utilized. Below, zooming to the nucleotide level shows that *D. yakuba* two mutations and a single nucleotide deletion relative to *D. melanogaster* have disrupted the canonical PAS (red box, asterisk), but that a less frequently utilized *D. melanogaster* PAS located downstream is converted into a dominantly used testis PAS in *D. yakuba* (blue box). This conserves the overall pattern of testis 3’ UTR shortening via different cis-regulatory sequences. (C, D) Screenshots of a *CrossBrowse* session loaded with small RNA-seq data from male bodies. (C) Examining orthologous miRNAs *dme-mir-960, dsi-mir-960*, and *dse_335* reveals a difference in the relative accumulation of the 5p vs 3p arms between *D. melanogaster, D. simulans, and D. sechellia*, respectively. (D) An indel occurring between homologous sequences in *D. pseudoobscura* and *D. persimilis* expresses miRNA *dps_3417*.

By zooming to the primary sequence, we observe that alteration of polyadenylation events at both *Pde1c* and *CG2201* are accompanied by mutations in a proximally located PAS. Notably, there exists an alignable proximal PAS in *D. yakuba CG2201* bearing only a single nucleotide change relative to its ortholog (**Figure 3A**). While one might have inferred this to represent a weak variant site, inspection in *CrossBrowse* makes it plainly evident that this altered PAS is essentially non-functional. On the other hand, the multiple changes to the *D. yakuba Pde1c* proximal PAS can more easily be imagined to render it inactive (**Figure 3B**). Interestingly, inspection of the primary sequence of the testis-specific polyadenylation event in *D. yakuba* reveals usage of a putatively cryptic PAS that is unmasked by loss of the upstream polyadenylation signal (**Figure 3B**), thereby preserving the pattern of testis 3’ UTR shortening. Notably, this signal is embedded within an identical sequence block in *D. melanogaster*, but does not yield any functional 3’ termini in this species.

Overall, these examples highlight different modes of functional divergence between otherwise very related species. Importantly, these evolutionary changes in functional elements would have been difficult to infer from primary sequence alone, and might otherwise require extensive bioinformatics to appreciate the diversity of evolutionarily changing patterns. Instead, they can be effectively uncovered and classified by human interaction with multiple types of cross-species deep sequencing data via *CrossBrowse*.

### Direct visualization of alterations in miRNA locus structure and processing

As another application in the post-transcriptional realm, we illustrate the utility of *CrossBrowse* for interpreting miRNA evolution. While miRNA loci were originally studied with respect to conserved arrangement and processing, presumably reflecting their orchestration of conserved target networks, the advent of deep sequencing of small RNAs across multiple species clades has permitted evolutionarily divergent aspects of miRNA pathway to be studied^22^-^24^.

We loaded *CrossBrowse* sessions with small RNA datasets from closely related groups of Drosophilids, namely *D. melanogaster/simulans/sechellia*, and *D. pseudoobscura/persimilis* (Mohammed and Flynt et al, in preparation), and browsed them for evidence of evolutionary lability in miRNA content and/or processing. Based on our recent appreciation of accelerated divergence in testis-restricted miRNA clusters^22^, we were interested to browse their behavior using our platform.

Candidate events of miRNA emergence can be categorized bioinformatically, but these require conscientious manual vetting as many fortuitous degradation products might masquerade as miRNAs^24^. However, it is extremely tedious using existing genome browsers to gain confidence that these represent genuine birth events in a specific species, and if so, what genomic alterations lead to their emergence. By simply inspecting the raw data in *CrossBrowse*, we could identify cases of “arm-switching” in the dominant accumulation of 5p vs 3p hairpin species within a a testis cluster in species of the melanogaster subclade (**Figure 3C**). This alteration is expected to alter the functional output of a miRNA locus^25^. More dramatically, we could also intuitively visualize a genomic alteration in a testis cluster that results in the recent birth of a miRNA locus within an individual species in the obscura subclade (**Figure 3D**).

Overall, such observations can be the basis of directed computational analyses, whose outputs can then be manually validated using *CrossBrowse*. Such cycles of computational and human assessment comprise an efficient workflow to interpret cross-species deep sequencing data.

### Conclusion: a generic tool for integrating cross-species datasets

The proliferation of public genomic experiments across many species enables opportunities for comparisons between species. Unfortunately, most existing browsers are difficult to configure, do not adequately visualize synteny and rearrangements between genomes, are not capable of displaying experimental data between species, and are not scalable to vertebrate-sized genomes. These limitations inhibit users, especially data producers who may not have substantial computational expertise, to quickly ask new questions using genomic datasets. We address these issues with *CrossBrowse*, an easily configurable comparative browser that supports standard genomics data formats and permits intuitive visualization of synteny and structural changes across the full gamut of genomic windows of interest. Data access routines are performed in background threads to maximize responsiveness of the user interface. This browser is implemented in Java, can be easily be installed and run on any platform with a JVM, and does not require internet access or data upload, allowing maximum flexibility for data with privacy requirements. We anticipate that *CrossBrowse* will find broad utilization, and help foster direct interrogation of cross-species data by bench scientists with limited computational expertise. Indeed, recent years have shown that several basic features of regulatory networks, RNA processing, and gene expression in the well-studied organisms are actively evolving, or have proven to be atypical or derived in some fashion. This highlights the importance of a broader cross-species foundation for experimental biology in the future.

## Methods

### Algorithm for constructing synteny representation

Whole genome alignments provide a means of mapping coordinates between two genome assemblies, from large regions down to nucleotide resolution. To generate an adaptive representation of these mapping functions, we down-sample the mapping function to render a visual representation of primary sequence alignment that is appropriate to the scale of the view. The synteny visualization is generated from a graph of the genomes represented in the session, where the genomes represent the nodes of the graph, and the whole genome alignment represent edges between two genomes. Since *CrossBrowse* represents synteny between neighboring pairs of genomes, the order in which they are stacked in the session induces an ordering between genomes. The top genome in the browser is labeled as the “source” genome, and the bottom genome is labeled as the “sink”. The source genome determines the parameters of the synteny visualization. The base-pair width of the segment displayed in this view determines the resolution of the syntenic blocks. The resolution is determined to be the maximum size such that at least the minimum width of a syntenic block in the source genome is 20 pixels. For example, if the browser window is 1000 pixels across, and displaying a region that is 100 bp long, 2 pixels will be the width allocated to each nucleotide, and one syntenic block will represent 10 nucleotides in the source genome.

The granularity of the source genome establishes interval partitions, whose register is aligned such that syntenic blocks line up with “round” coordinates in the source genome (1, 5, 10, 25, 50…). This partitioning establishes the “base” of each syntenic block. Syntenic blocks are constructed using a recursive algorithm that is applied to each interval in the source genome. For each interval, whole genome alignments are queried to project the coordinates of source sequence into the coordinate system of the next genome in the stack. The source coordinates are added to a stack, and the coordinates in the target genome become the new source interval. This process is repeated until coordinate translation either reaches the “sink” genome, or there are no remaining segments for which a valid coordinate translation exists.

There are a number of edge cases that must be considered when performing these projections. For a given query interval, the whole genome alignment may not allow coordinate translation for a subsequence (or entirety) of the queried segment. If these gaps occur on the boundary of a syntenic block, this will have the effect of “narrowing” the syntenic block, and must be propagated along the sequence of preceding syntenic segments to faithfully represent the boundaries segments that are syntenic across all displayed genomes. Additionally, a single segment can map to multiple locations in a target genome. In this manner, the nascent syntenic blocks form a tree, where the source interval forms the parent node, which will projected onto one or more “child” segments in the target genome.

Once the syntenic intervals composing each syntenic block are assembled, graphical parameters are from the GUI, including the relative vertical space allocated to each genome, are integrated with parameters of the view indicating whether each genome should be drawn in a left-to-right or inverted coordinate space, to establish the coordinates used to render each synteny block. Additional genomes can be stacked if desired, and their relationship will be visualized by the above logic.

### Deriving CHAIN files to support rapid coordinate translation

*CrossBrowse* represents whole genome alignments using uses CHAIN files, which users can easily access from the UCSC genome browser. CHAIN files have a polarity, allowing for coordinate queries from a source genome to be translated into the coordinate space of a target genome. To enable facile exploration of multiple genomes simultaneously, *CrossBrowse* derives CHAIN files with the necessary polarity to support rapid queries in either direction between any pair of genomes with no additional user interaction. Additionally, whole genome alignments are often generated with a “star” topology (i.e. all organisms versus human), such that it is necessary to make multiple queries each time coordinates are between two species.

Represented as a graph where the genomes represent the nodes and whole genome alignments the edges between nodes, *CrossBrowse* includes the functionality to derive missing edges. To do this requires two fundamental operations, CHAIN inversion and CHAIN composition. Inversion is trivial to implement, and is essentially achieved by swapping the ‘source’ and ‘target’ fields of the chain, and ordering the chains with respect to the target genome. Given genomes A, B, and C, the composition *f□* ***1*** *g*, takes two input alignments f: (A→B) and g: (B→C) and outputs the mapping from (A→C). Addition is implemented by taking the boundaries of each aligned block in the source genome, and recursively translating its coordinates into the space of the ‘sink’ genome.

With these two fundamental operations in hand, we use a greedy algorithm to derive missing edges from a graph of genomes and their alignments. To begin, all available chains are inverted. Second, all pairs of nodes (genomes) are enumerated, and pairs for which no alignment exists are identified. For each pair without an alignment we identify the shortest path between those two nodes using existing edges in the graph, and tabulate the frequency of all 3-node subsequences (triplets) in the set of shortest paths. Taking the triplet that is common to the most missing paths, we apply composition on the two CHAIN files that form the edges between the three nodes, and the composition is inverted to generate the reverse edge. The length of all paths that contain this triplet reduced by one by the composition operation. This procedure is recursively repeated applied there are no nodes that have a path length of greater than two between them.

### Indexing CHAIN alignments

The UCSC CHAIN format is used to represent whole genome alignments. However, in its original form, the CHAIN format is not amenable to random access queries. TABIX indexing can be used to index many commonly used data formats. To enable TABIX indexing CrossBrowse converts conventional CHAIN format files to a row-oriented GFF format file. The resulting GFF file is compressed, indexed, and subsequently queried using the TABIX API provided by HTSJDK.

### Analysis of chromatin features

We downloaded human/chimp ChIP-seq data for chromatin features from Gene Expression Omnibus (GEO) accession GSE70751^13^, and CTCF ChIP-seq data from *Drosophila* species from GSE24449^16^. We also utilized CTCF peak calls annotated in *D. melanogaster* and *D.* pseuodoobscura, respectively (in files GSM602326_dmel_peak_regions.bed and GSM602335_dpse_peak_regions.bed).

We compared two strategies for determining the presence of orthologous CTCF binding sites. The first approach replicated the previously reported scheme^16^, in which we used *liftOver* to translate the coordinates of the 2,182 *D. melanogaster* peak calls to the *D. pseuodoobscura* genome. Sites that could not be translated were marked as *melanogaster-*specific. We intersected the remaining sites with the 2,332 CTCF binding intervals called in *D. pseudoobscura*, to annotated the overlap as conserved CTCF sites.

In the second strategy, we created two new intervals for each *D. melanogaster* CTCF peak call by alternately extending the original interval by 500 nt on either side. The coordinates of each extended interval were translated between assemblies. If a pair of intervals was (1) translated to the same reference sequence, (2) spanned a genomic interval no more than two-fold longer than the original *melanogaster* interval, and (3) in a relative orientation between the two was consistent with that observed for *melanogaster,* then they were considered to represent orthologous regions. As noted, this procedure rescues >200 *D. melanogaster* CTCF sites from being unaligned in *D. pseuodoobscura*, to having orthologous genomic assignments. We assessed these for experimental evidence of CTCF binding in both species, paying especial attention to manually assess all “rescued” conserved sites.

### Software Availability

CrossBrowse was implemented in Java. Executables and source code are available from https://github.com/shenkers/ComparativeBrowser/releases. An online document with detailed instructions for configuration and use are available at https://github.com/shenkers/ComparativeBrowser/wiki.

## Acknowledgements

We thank Piero Sanfilippo for access to 3’-end sequencing data, Alex Flynt and Jaaved Mohammed for observations on miRNA evolution, and Brian Joseph for manuscript comments. S.S. was supported by the Tri-Institutional Training Program in Computational Biology and Medicine. Work in E.C.L.’s group was supported by the National Institutes of Health (R01-NS074037, R01-NS083833 and R01-GM083300) and MSK Core Grant P30-CA008748.

## System requirements

The minimum requirement for running `CrossBrowse` is a system capable of running a Java 8 runtime environment. If you do not have Java installed on your computer it can be downloaded from https://java.com/download

## Installing CrossBrowse

The most up-to-date release of `CrossBrowse` can be downloaded from https://github.com/shenkers/CrossBrowse/releases/latest

The executable **JAR** file will have a name `CrossBrowse-<version>.jar. CrossBrowse` can also be installed by building the **JAR** from source. In addition to a Java development kit, you will need to install **Maven** and **git**. The source can be downloaded using the sequence of commands

~~~
**git clone https://github.com/shenkers/CrossBrowse.git cd CrossBrowse
mvn package**
The compiled JAR file can be found at **CrossBrowse/target/CrossBrowse-<version>.jar** relative to the folder where git clone was called.
~~~

## Running CrossBrowse

The `CrossBrowse` executable can be run using the following command:

**java -Xmx2G -jar CrossBrowse-<version>.jar**

This command will launch `CrossBrowse` with **2GB** memory via the `-Xmx` argument. If your `CrossBrowse` session is terminated by an `OutOfMemoryError`, the `-Xmx` parameter should be increased to provide **CrossBrowse** with more memory.

## CrossBrowse configuration

When the browser is first started you will be presented with an empty window as follows:

The first step to configure the genome browser is to identify the genomes you will be using. To do this select **`File -> Load Genome`**.

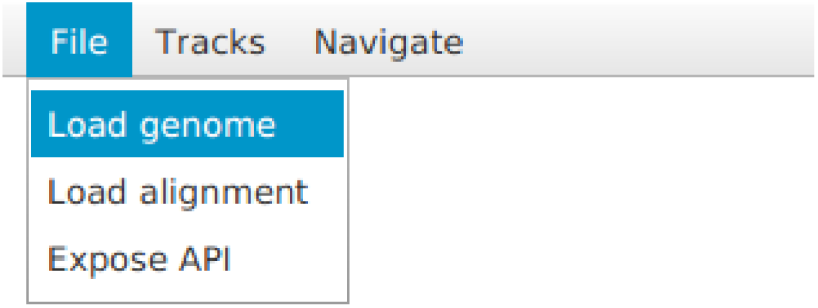

You will be presented with the following prompt:

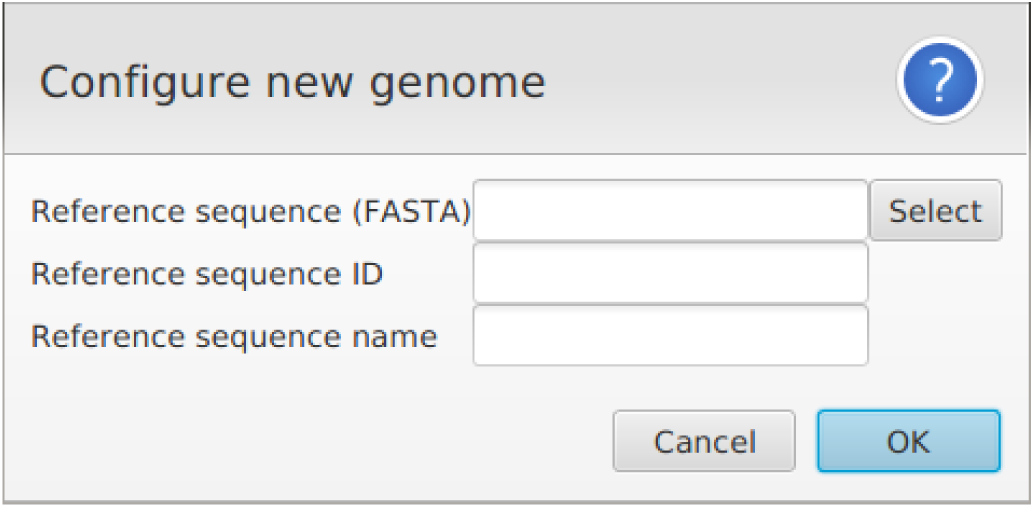

where you can specify the identifier for the genome you will be using. To view multiple genomes in a single session simply use this menu to add additional genomes. Once you click Ok the main view window will be updated to include a new partition for the added genome, as shown below (here 3 genomes have been loaded):

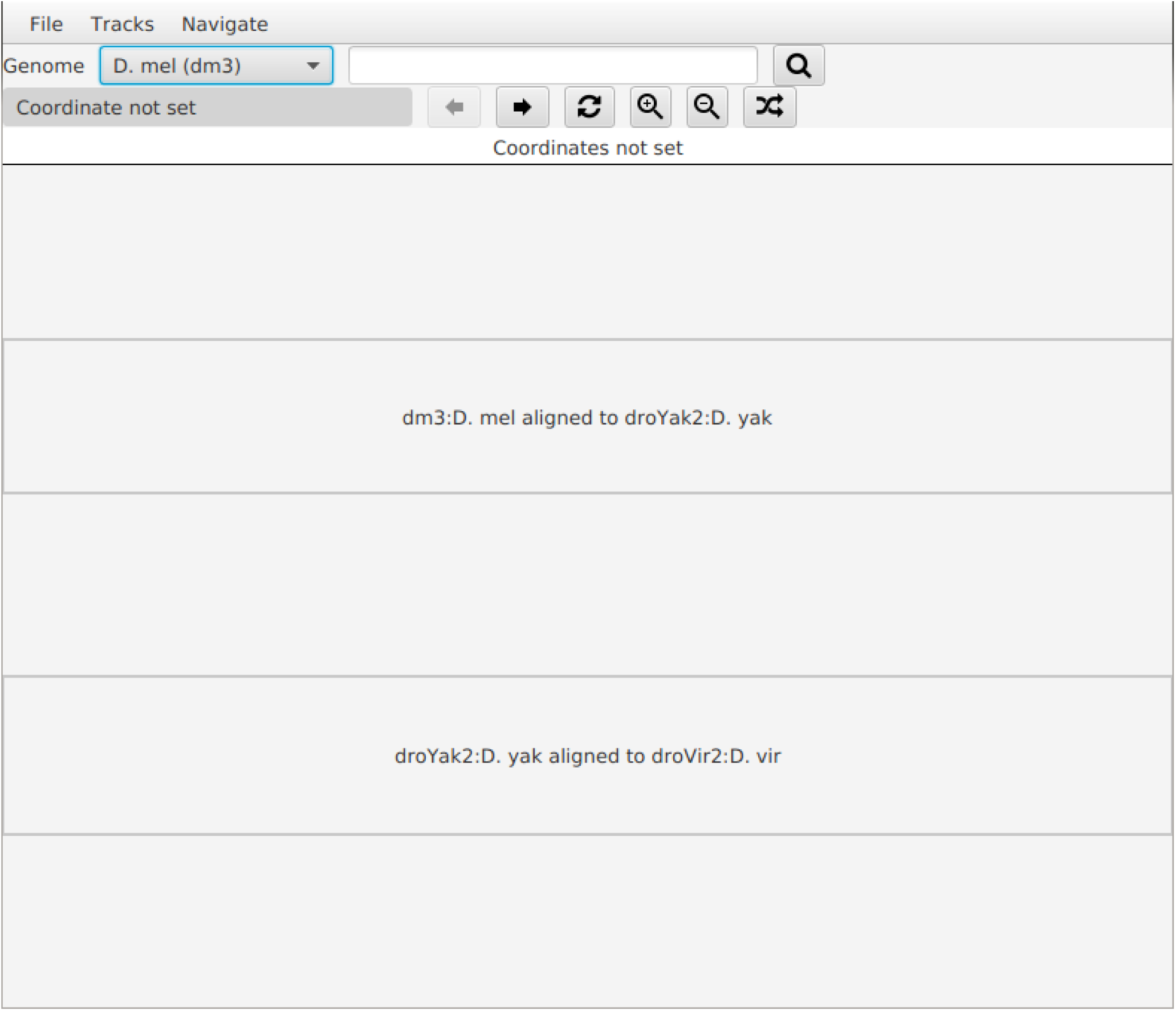

### Loading tracks

Once at least one genome has been specified, you can start loading data tracks by going to the menu and selecting **`Tracks -> Load Track`**.

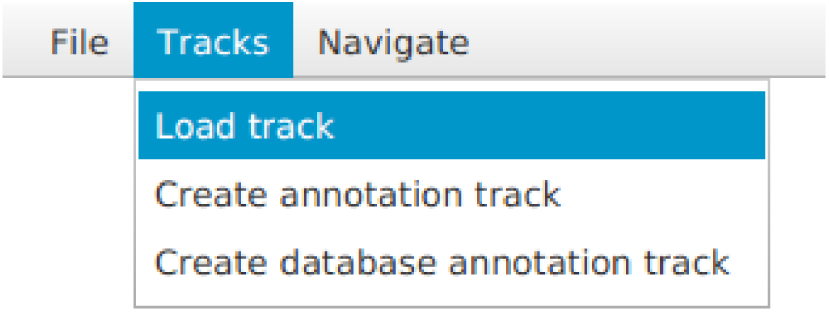

You will be presented with the following prompt:

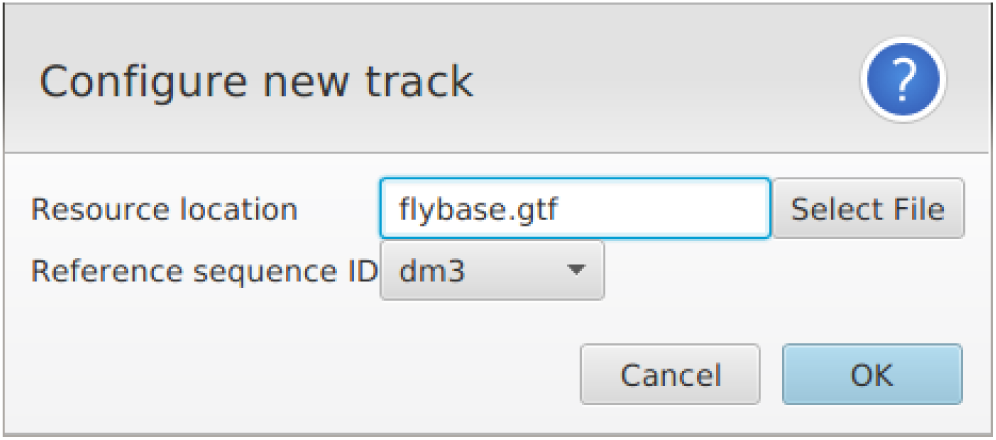

Where you can select the files you wish to load, and the genome that they should be displayed relative to. Using this menu the browser can load BAM, BED, BW (big-wig), GTF format files. After loading some tracks for each genome and setting the viewed coordinates, the browser should look something like this:

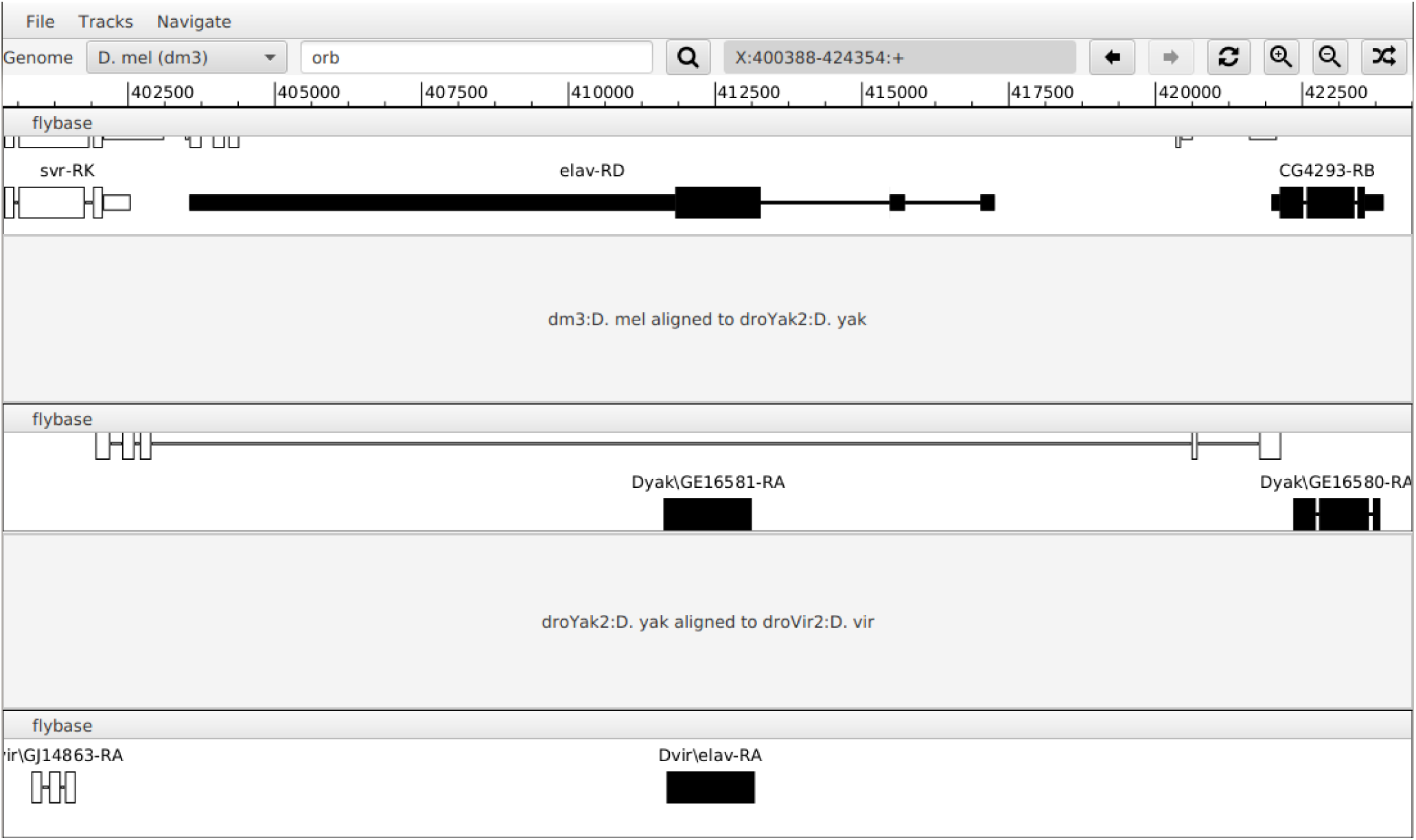

To enable efficient access, some flat files (BED, GTF) are indexed before loading, so you may experience a delay the first time the file is loaded by the browser. The generated index is cached on disk, so these delays are avoided in subsequent sessions.

## Loading genome alignments

Once all your genomes have been loaded you can load whole genome alignments by going to the menu and selecting **`File->Load Alignments`**.

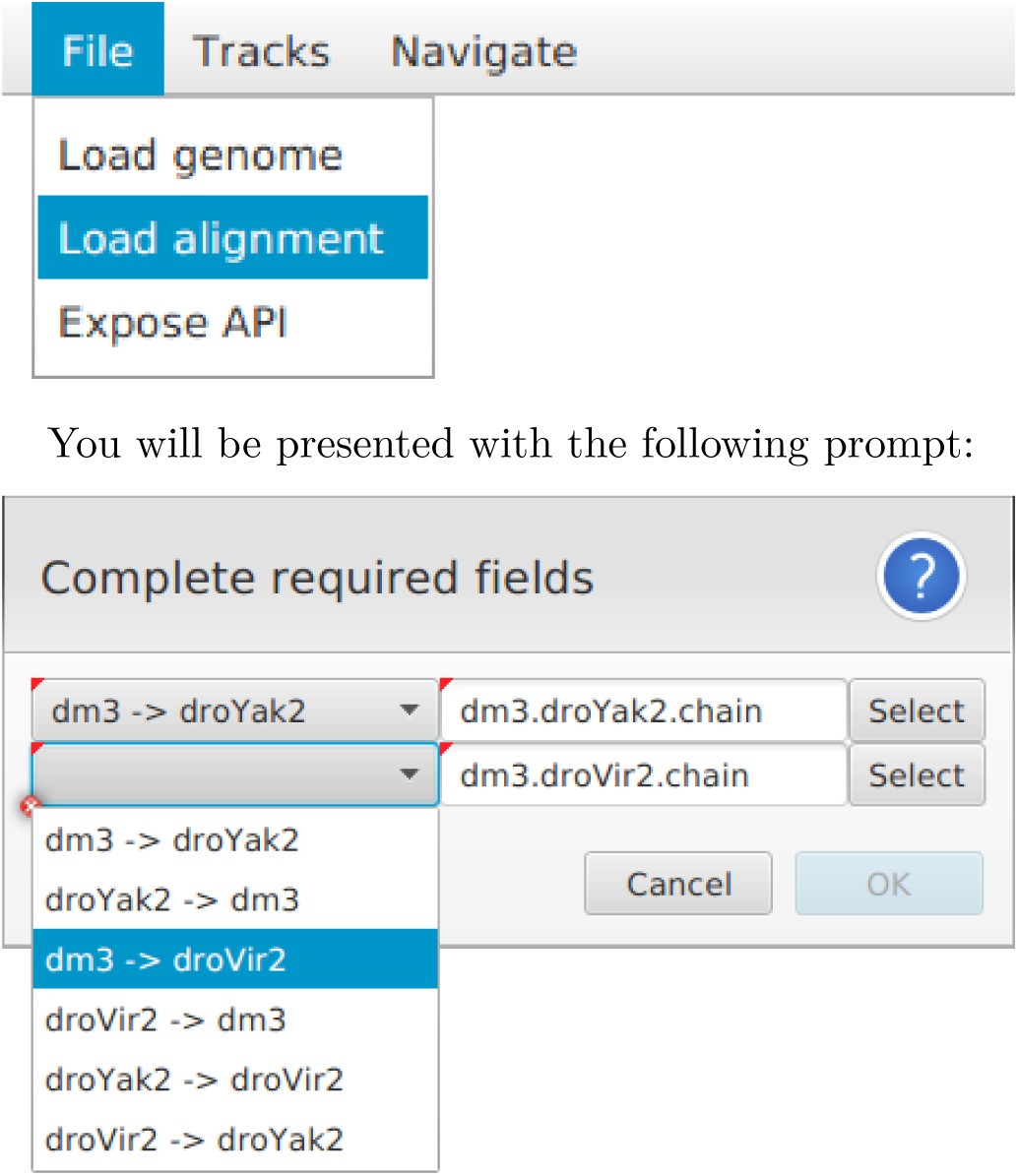

You will be presented with the following prompt:

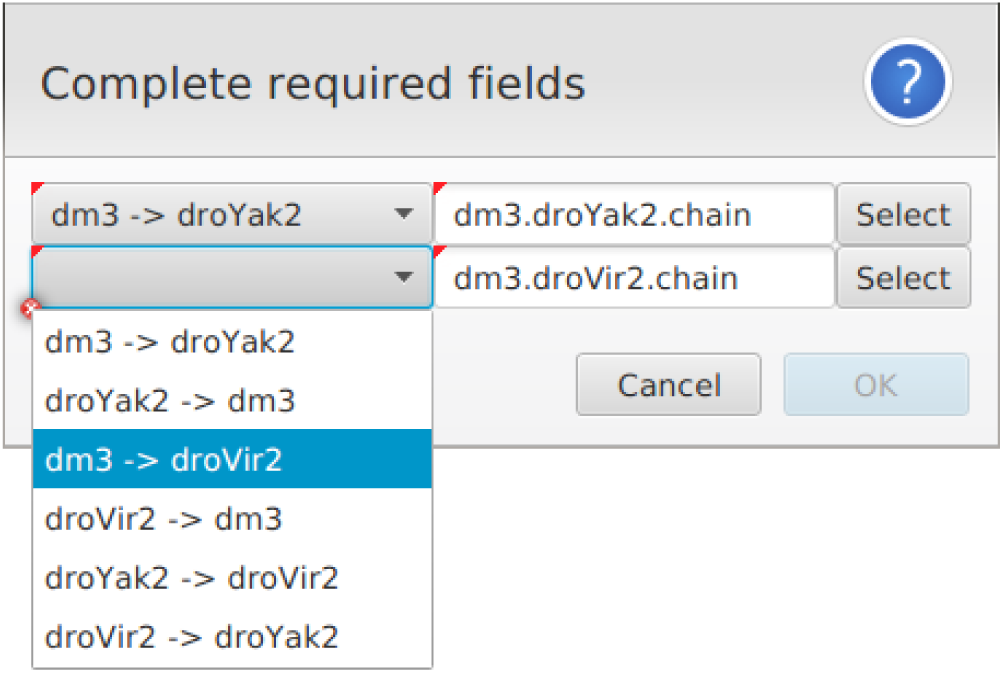

that allows you to load pairwise alignments between the genomes in your session. Here, if the session has k genomes loaded, it is only necessary to load k-1 pairwise alignments such that all your genomes are mutually reachable. Consider the case where k=3, then there are 3 valid combinations of alignment files you can load at this window as depicted below.

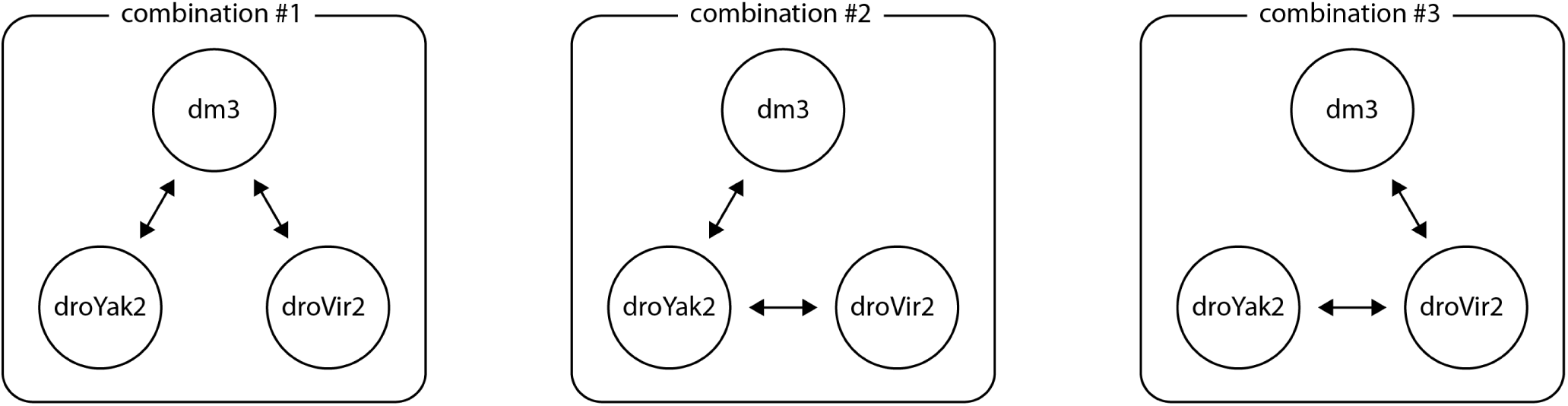

Once the alignments are loaded the synteny overlay will be displayed:

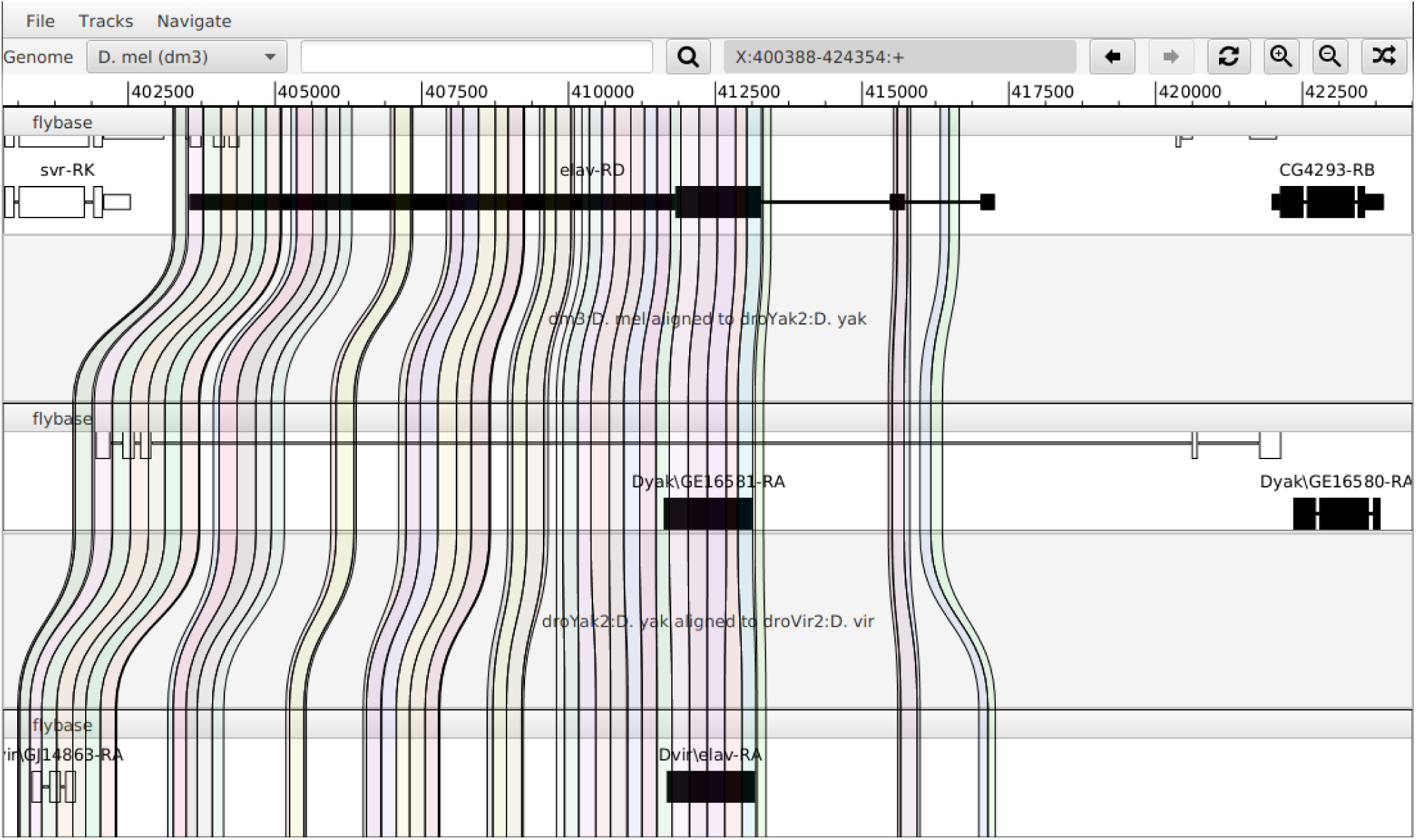

## Navigation

The browser has familiar navigation interface. At the top of the window there is a navigation bar that allows you to enter a specific genome coordinates. The genome to adjust can be selected from the drop down menu. Similar to the UCSC genome browser, the first textbox allows entry of gene by name or specific coordinates. A second text box displays the coordinates of the currently displayed window.

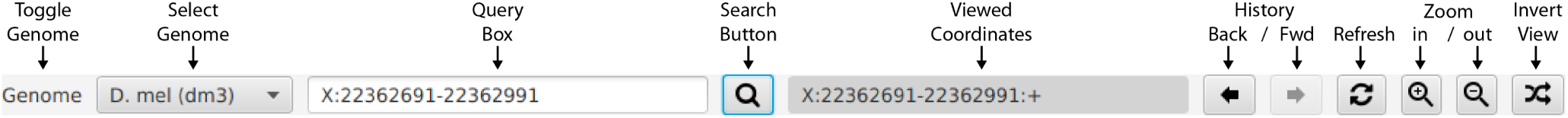

Using the query box you can set the viewed coordinates for each genome by specifying an explicit range to be viewed (as depicted above) or by searching for features in loaded annotation files. For example, to search by gene-name, type the gene name into this box, and the browser will suggest matching features as shown below:

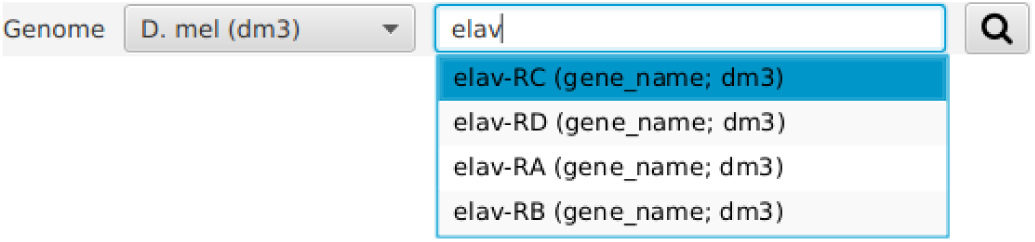

Using the mouse one can re-center the viewed window by clicking on a single position, as below:

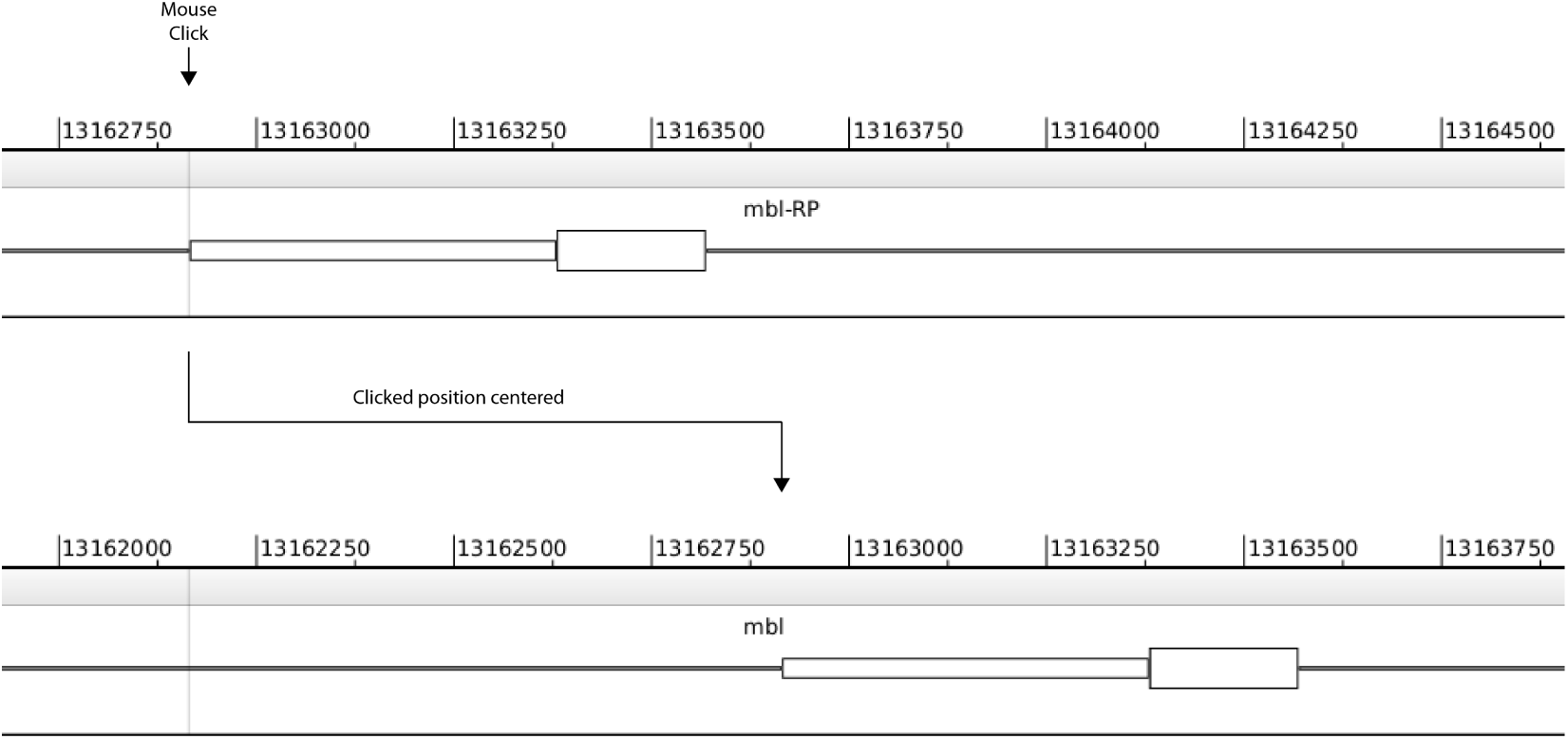

Similar to existing genome browsers, you can zoom in on a slice of the current view by using a mouse drag gesture to select the coordinates you would like to zoom in on:

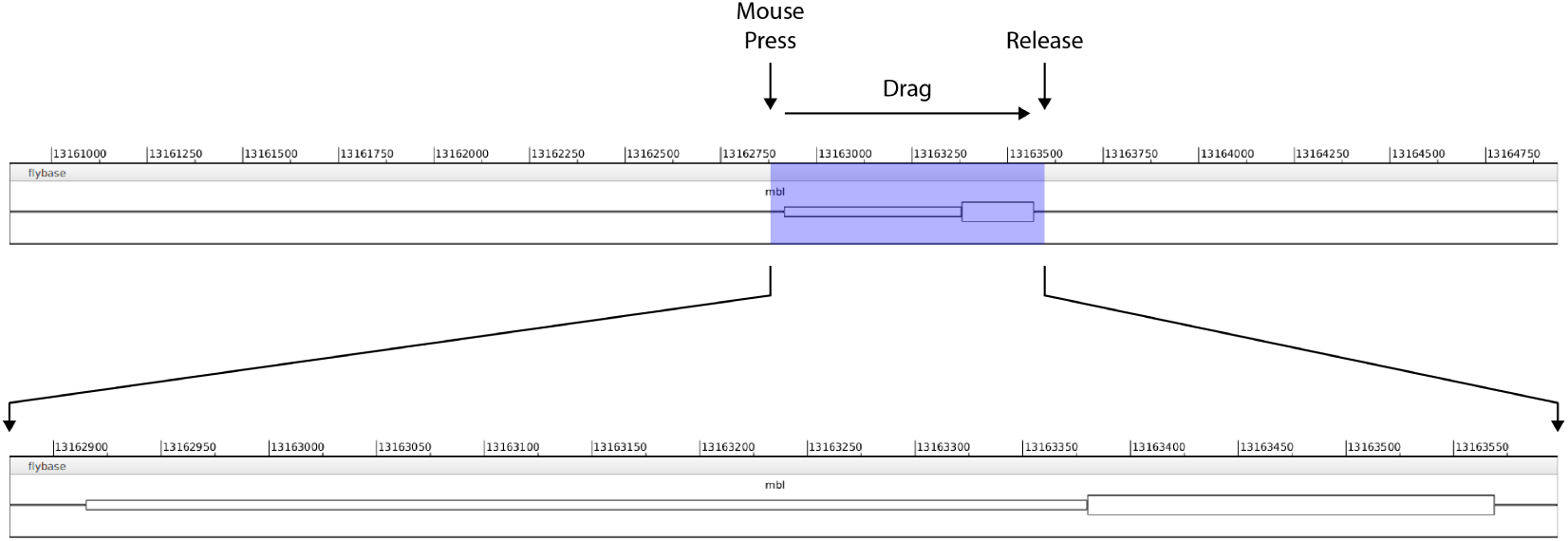

Zoom buttons 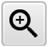 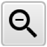 allow the displayed segment to be zoomed in and out:

In some instances the orientation of gene is flipped between two Genomes. In this case, the 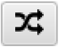 button can be used to invert the frame of reference, so that chromosomal coordinates are displayed in decreasing order from left-to-right.

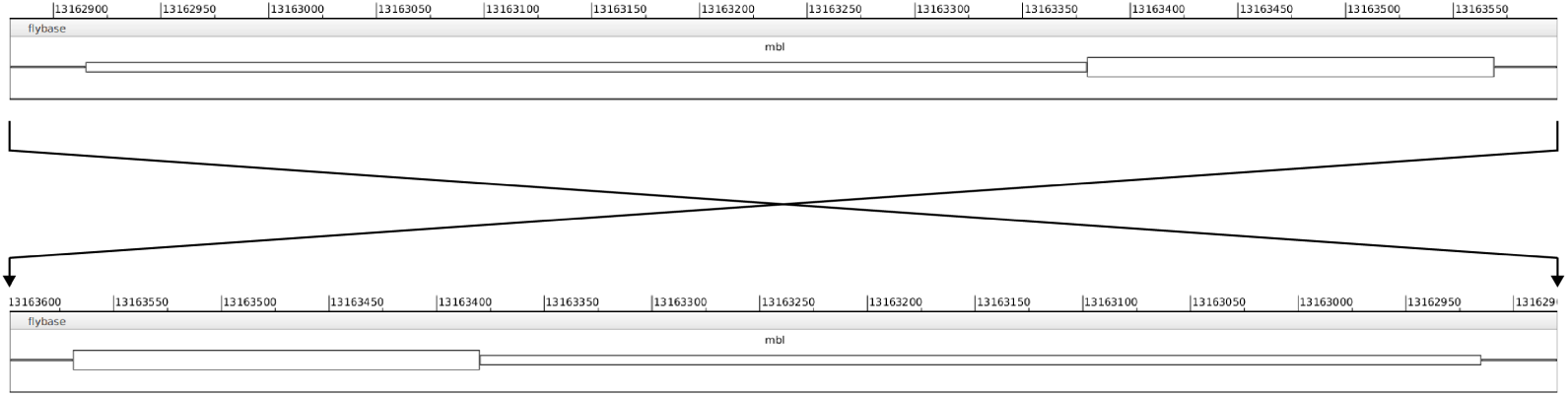

### Coordinate translation (liftOver)

Once pairwise alignment have been loaded the liftOver functionality will be enabled from the **Navigation** menu:

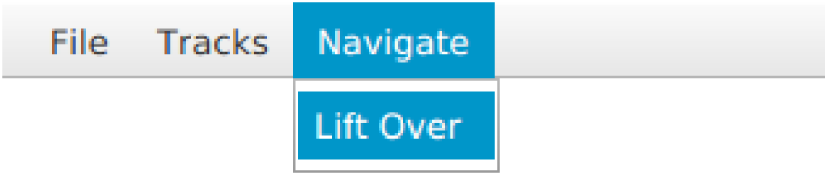

You will be presented with the following prompt:

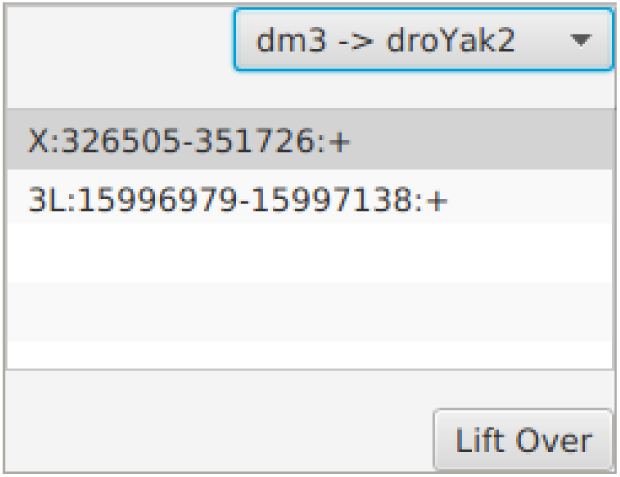

The toggle box in the upper right hand corner of this window allows the **source** and **target** genomes to be used for coordinate translation. In the above example we have selected **`dm3 -> droYak2`** which means that liftOver will use the alignment and the coordinates of the view for the **`dm3`** genome to set the coordinates in **`droYak2`** genome. The list at the center of this window lists all segments in the **`target`** genome that have an alignment with the interval displayed for the **`source`** genome. In this case the viewed segment of the **`dm3`** genome maps to two different contigs in the **`droYak2`** genome. Selecting one of these locations and clicking the **`liftOver`** button at the bottom of this window will set the displayed coordinates in the **target** genome accordingly. If you leave the **`liftOver`** panel open, it is dynamically updated as you navigate so that you rapidly access syntenic coordinates in all displayed genomes.

## Annotation tracks

In addition to loading data tracks, **`CrossBrowse`** includes special facility for for creating annotations while browsing. To create a new set of annotations select **`Track -> Create annotation track`** from the main menu.

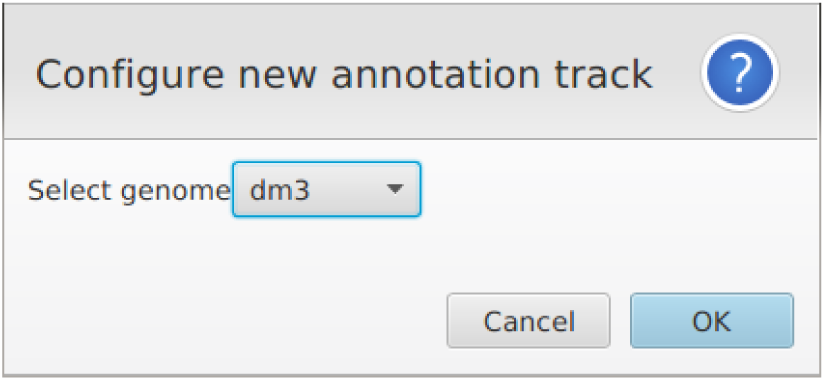

You will be prompted to enter specify the species with which it should be associated. Once the track is created, mouse gestures can be used to add new annotations to this track while you are browsing. A single click will create a single-nucleotide interval. Using a click-drag-release gesture will create a stranded annotation, where the initial position that is clicked is the 5’-end of the interval.

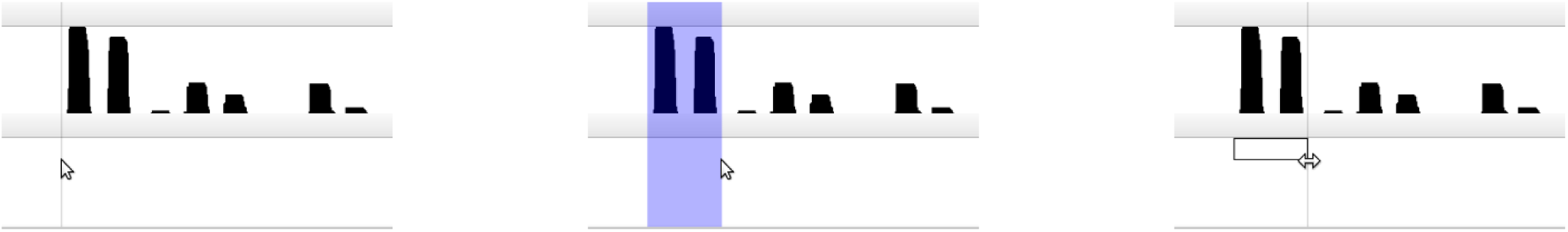

If you would like to adjust the coordinates of an annotation, clicking and dragging the boundaries to make the necessary adjustment.

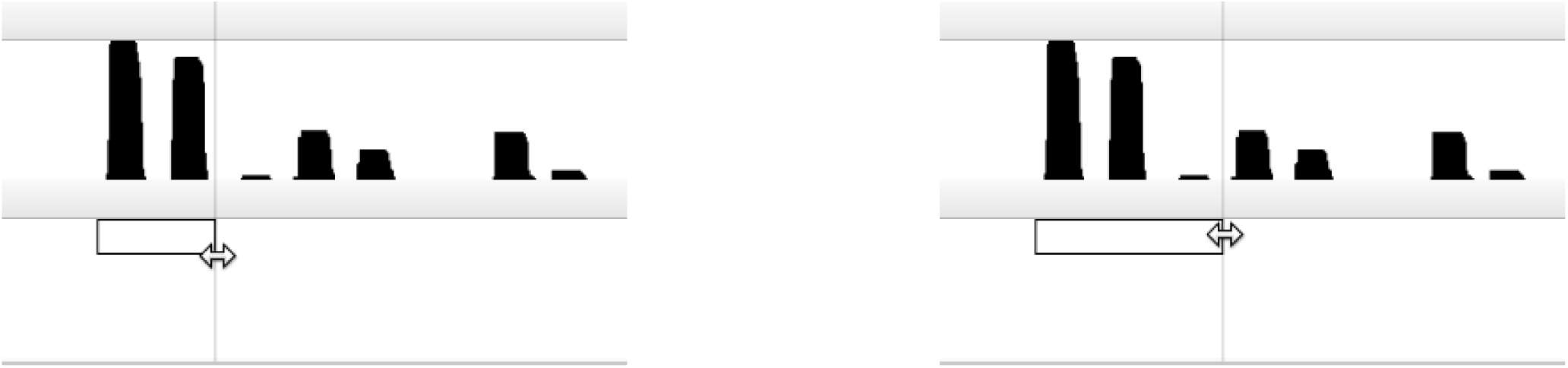

Right clicking on an annotation interval will bring up a context-menu enabling the strand assigned to the interval to be changed, or for the annotation to be removed entirely.

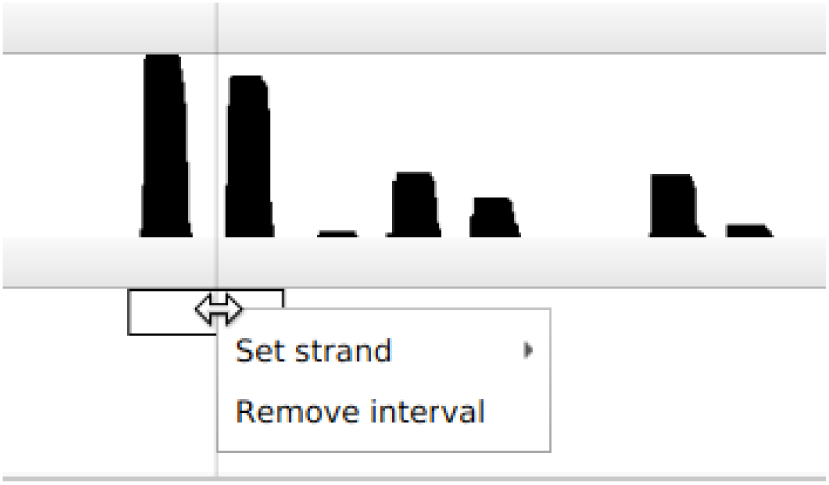

Right clicking on an empty position in the annotation track will bring up another context menu allowing you to import or export intervals to or from the track from and BED format files.

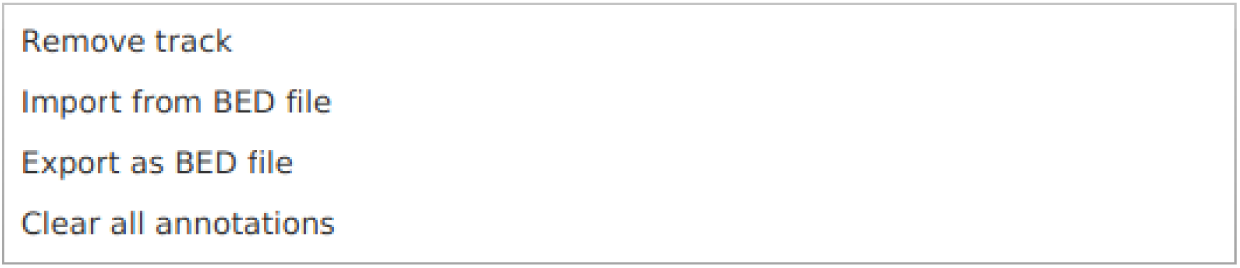

In addition to regular annotation tracks, an option to create a database-backed annotation track is also available. The lifespan of a regular annotation track is a single CrossBrowse session. It is the user’s responsibility to manage saving work when finished, and managing those annotations between sessions. The database backed annotation track provides the same functionality as a regular annotation track, and automatically saves the annotations to a database as they are created. In this manner, the user can access a set of annotations over a series of **`CrossBrowse`** sessions, and will not lose any annotations if their session is interrupted before saving any updates. To create a database annotation track select **`Track -> Create database annotation track`** from the main menu.

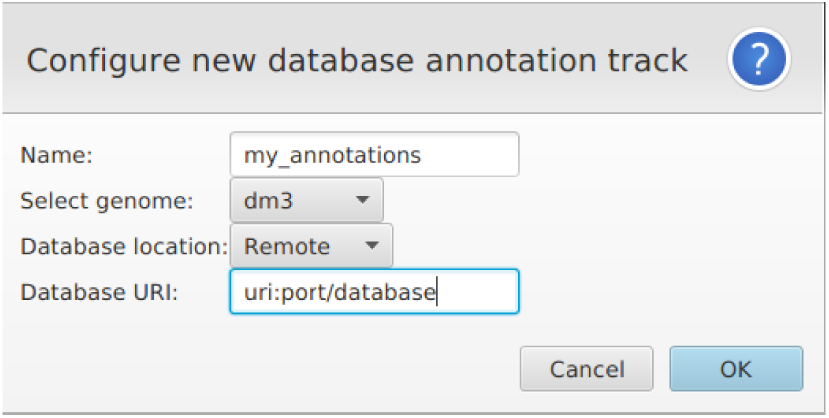

A prompt will allow you specify the location of the database. A local database will be written directly to your computer. Alternately, the track can be backed by a remote database. In this case, a URL and port will be used to specify the location of the database.

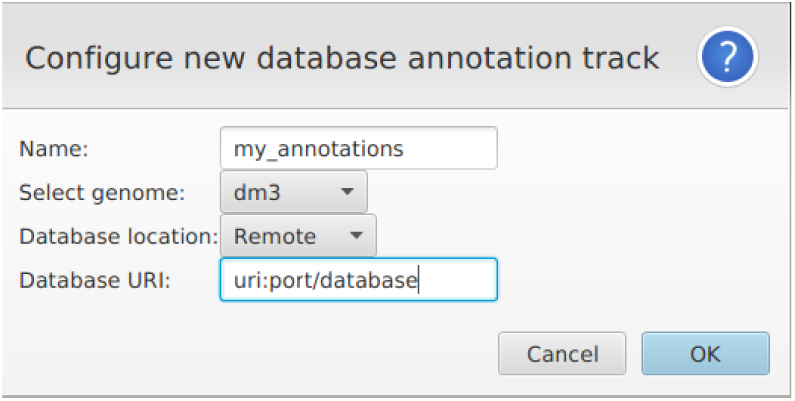

At the current time, the remote database must be Derby database.

## Programmatic control of CrossBrowse

`CrossBrowse` provides a web API that enables aspects configuration and navigation to be controlled automatically. To access this functionality launch the web-server by selecting **`File -> Expose API`**

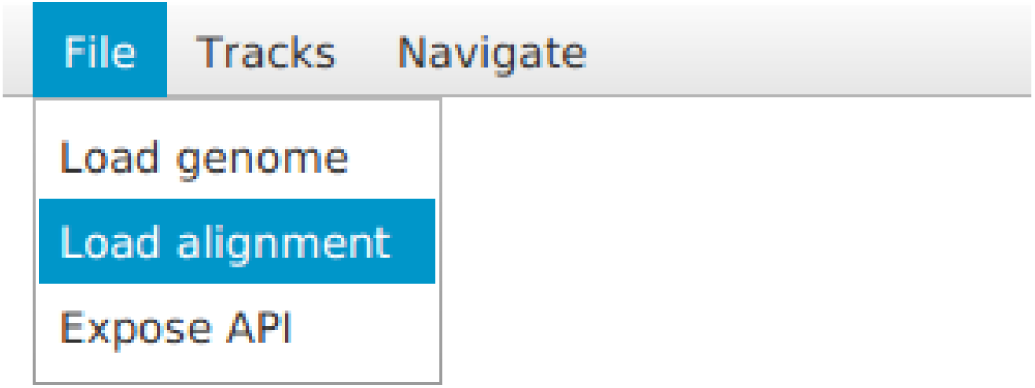

This will display a prompt asking for a port number. Entering a port number and pressing Ok will launch the **`CrossBrowse`** web server listening at URI **http://localhost:port**. The web API can be used to query and set displayed coordinates, load tracks, and take snapshots of the current session. The basic structure of a web request is:

~~~
[URI]/[command]?[param-1]=[value-1]&[param-2]=[value-2]…[param-n]=[value-n]
~~~

To experiment with the API you can perform requests using the URLs in a web browser, or command line utilities such as **curl** or **wget** capable of making web requests. The available commands and their parameters are described below.

## Commands

### Coordinates

- getCoordinate (HTTP GET) ~~~
– parameters
           * genomeId (String) -id of the genome in the current session. Optional, if omitted coordinates of the id of the first genome will be used.
– returns
          * A plain test string of the form ‘chr:start-end:toNegativeStrand’
– example
          * localhost:12345/getCoordinates
~~~

- setCoordinates (HTTP GET) ~~~
– parameters
          * genomeId (String) -id of the genome in the current session. Optional, the id of the first genome will be used if omitted
          * chr (String) -the name of the chromosome
          * start (int) - the start coordinate of the requested view
          * end (int) - the end coordinate of the requested view
          * toNegativeStrand (boolean) - whether the view should be inverted
– returns
          * A HTTP OK response on sucess
– example
          * localhost:12345/setCoordinates?chr=X&start=100&end=200&toNegativeStrand=false
~~~

### Tracks

- loadTrack (HTTP GET) ~~~
– parameters
         * genomeId (String) - id of the genome in the current session. Optional, the id of the first genome will be used if omitted
         * uri (String) - path to the resource to be loaded
– returns
         * A HTTP OK response on sucess
– example
         * localhost:12345/loadTrack?genomeId=dm3&uri=/path/to/file.gtf
~~~

### Snapshot

- getSnapshot (HTTP GET) ~~~
– parameters
           * scale (float) - Image sampling resolution used for the snapshot. Optional, if omitted a default scale of 1.0 (screen resolution) will be used.
           * tracksOnly (boolean) - whether the menu bar should be included in the snapshot
– returns
           * A PNG image of the CrossBrowse session
– example
           * localhost:12345/getSnapshot?scale=2.0&tracksOnly=false
~~~

**Getting help**

Online documentation can be found on the **`CrossBrowse`** wiki https://github.com/shenkers/CrossBrowse/wiki. If you have a problem using CrossBrowse please create an issue at https://github.com/shenkers/CrossBrowse/issues.

